# scAllele: a versatile tool for the detection and analysis of variants in scRNA-seq

**DOI:** 10.1101/2022.03.29.486330

**Authors:** Giovanni Quinones Valdez, Ting Fu, Tracey Chan, Xinshu (Grace) Xiao

## Abstract

Single-cell RNA sequencing (scRNA-seq) data contain rich information at the gene, transcript, and nucleotide levels. Most analyses of scRNA-seq have focused on gene expression profiles, and it remains challenging to extract nucleotide variants and isoform-specific information. Here, we present scAllele, an integrative approach that detects single nucleotide variants, insertions, deletions, and their allelic linkage with splicing patterns in scRNA-seq. We demonstrate that scAllele achieves better performance in identifying nucleotide variants than other commonly used tools. In addition, the read-specific variant calls by scAllele enables allele-specific splicing analysis, a unique feature not afforded by other methods. Applied to a lung cancer scRNA-seq data set, scAllele identified variants with strong allelic linkage to alternative splicing, some of which being cancer-specific. scAllele represents a versatile tool to uncover multi-layer information and novel biological insights from scRNA-seq data.

## Introduction

Single-cell RNA sequencing (scRNA-seq) affords a unique glimpse into the transcriptome at the single-cell resolution, revealing great cellular heterogeneity *(1)*. Although this type of data harbors rich information of a cell’s transcriptome, most studies focused exclusively on gene expression, without tackling other important aspects such as single-nucleotide variants (SNVs) *(2)* or allele-specific expression *(3*–*5)*. In addition, previous analyses of genetic variants in scRNA-seq data used methods originally designed for bulk RNA-seq or DNA-seq *(6, 7)*, due to the lack of tools specifically tailored for variant calls in scRNA-seq.

Variant calling in RNA poses significant computational challenges. Typically, variant callers rely on resolving the haplotypes from the NGS reads *(8, 9)*. However, this strategy has limited applications at the RNA level where alternative splicing, allele-specific expression or RNA editing may affect minor allele frequencies of the variants or obscure the true haplotype proportion in the reads. Furthermore, in scRNA-seq, most expressed genes have shallow coverage *(10, 11)*, highlighting the need of accurate variant detection with limited reads. To date, no method exists that explicitly focuses on both SNVs and insertions/deletions (INDELs) in scRNA-seq.

Following their identification, the next major challenge is to link nucleotide variants to their potential molecular function. To this end, RNA-seq data possess unique advantages given the afforded multi-level information: gene expression, transcript isoforms and sequence variations. Using bulk RNA-seq, numerous studies leveraged this strength to uncover allelic bias of genetic variants in gene expression or splicing *(12, 13)*, which, for example, can lead to discovery of functional *cis*-acting variants that alter splicing *(13*–*16)*. Despite their typical low coverage, scRNA-seq data provide similar multi-level information. However, no method exists to exploit such features of scRNA-seq to uncover functionally relevant variants.

Here, we introduce scAllele, a versatile tool that performs both variant calling and functional analysis of the variants in alternative splicing using scRNA-seq. As a variant caller, scAllele reliably identifies SNVs and microindels (less than 20 bases) with low coverage. It implements RNA-friendly haplotype filtering by accounting for potential RNA editing sites and allele-specific splicing. Following variant calling, scAllele identifies significant associations between variant alleles and alternative splicing, which provides direct evidence of allele-specific splicing.

Using scRNA-seq data associated with well-characterized genotypes, we show that scAllele outperforms other commonly used methods, especially for microindel identification. We apply scAllele to scRNA-seq data derived from lung cancer samples. Our analysis identifies variants that have significant allelic linkage to splicing isoforms, some of which are enriched in cancer cells. Thus, scAllele is an integrative analysis tool that uncovers multi-level information in scRNA-seq.

## Results

### Algorithm Overview

scAllele calls nucleotide variants via local reassembly (Fig. 1a). To scan variants in the entire transcriptome, we split the mapped reads into read clusters (RC), defined as genomic intervals containing overlapping reads. Reads from each RC are subsequently decomposed into overlapping k-mers and reassembled into a directed de-Bruijn graph. The reference genomic sequence is included in the reassembly to serve as the reference haplotype in the RC. The nodes of the graph represent k-mers derived from the read sequence. Two nodes with k-mers overlapping by k-1 bases are connected with a directed edge. The ‘bubbles’ in the graph represent differences among all sequences including the reads and genome reference sequence.

**Figure 1.**
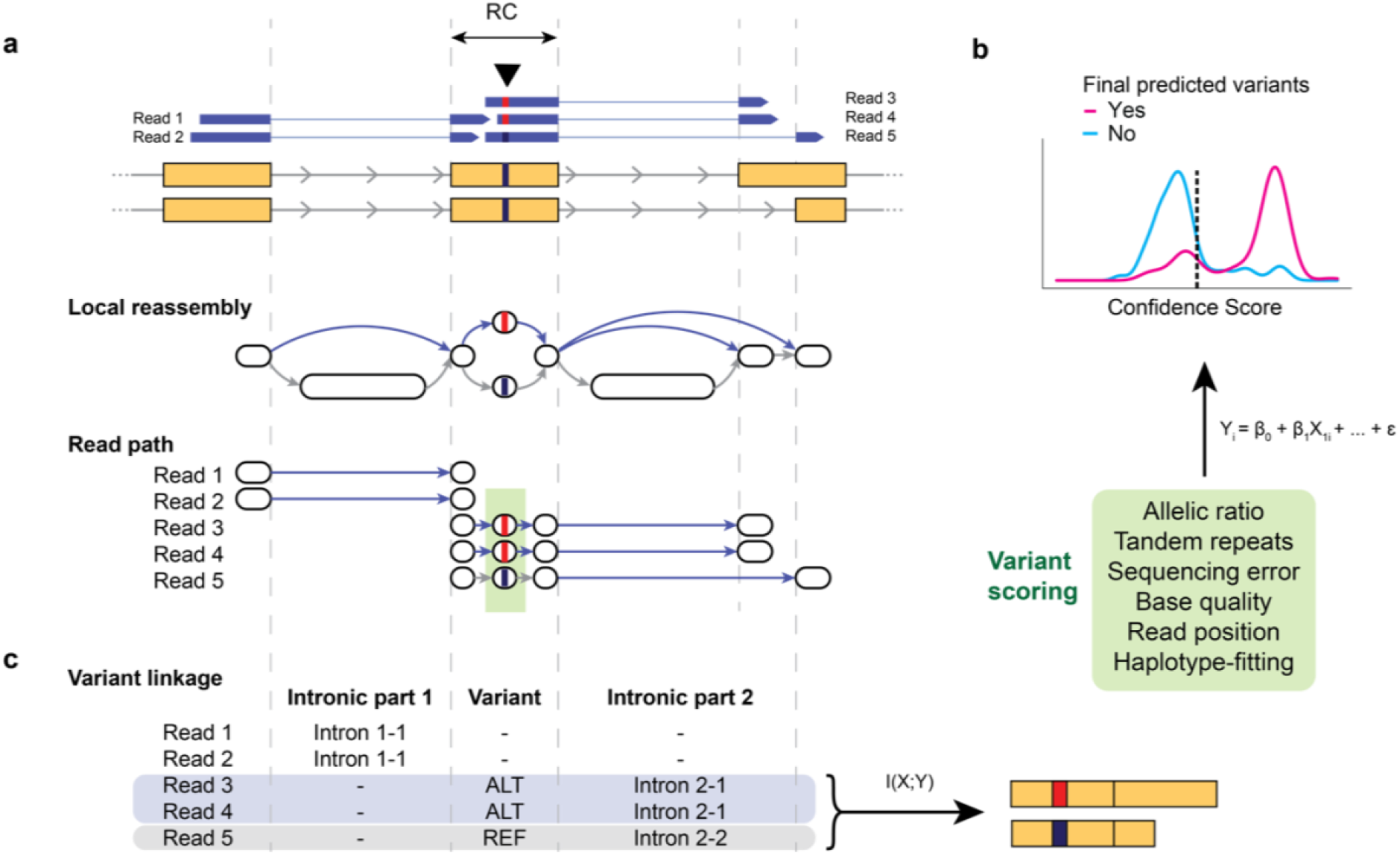
Algorithm outline. a. Illustration of the main algorithm of scAllele for variant calling. The reads and the reference genomic sequence overlapping a read cluster (RC) are decomposed into k-mers and reassembled into a de Bruijn graph. The graph shown here is a compacted version. The ‘bubbles’ in the graph indicate a sequence mismatch i.e., a variant. For each read, scAllele obtains a path for the original read sequence and infers the allele of each variant (including introns). b. Variants (green box in a) identified from the graph are then scored using a generalized linear model (GLM). The GLM was trained with different features (green box) to assign a confidence score to the variants. See Methods for details. c. To identify allele-specific splicing (i.e., variant linkage), scAllele performs a mutual information calculation between nucleotide variants (SNVs, microindels) and intronic parts (where the ‘alleles’ are the different overlapping introns), to calculate allelic linkage of splicing isoforms.

To identify nucleotide variants, we first traverse the graph with a depth-first search to identify nodes marking the beginning and the end of each bubble (source and sink nodes) and their respective pairing (Fig. 1a, Local reassembly). Hereafter, we perform a per-read analysis of the graph, where we first obtain the walk in the graph that best matches the read sequence, followed by the identification of variants present in each read. The presence of repeats or low-complexity regions significantly complicates the detection of variants since the de-Bruijn graph can be traversed in multiple ways. scAllele overcomes this challenge by performing a Dijkstra-based traversal of the graph with the assumption that the walk with the smallest editing distance best represents the set of variants present in the read. Finally, we collect the variants from the RC reads and score them using a generalized linear model (GLM) (Fig. 1b) where the following features are included: read position, base quality, number of neighboring tandem repeats, allelic ratio, sequencing error rate, and haplotype fitting (Methods).

An important feature of scAllele is the detection of variants at the read level. This feature enables a direct analysis of allelic linkage between the variants and other attributes of the reads. Here, we focus on identifying allelic linkage with alternative splicing via mutual information (Methods), similarly as in our previous work for RNA editing identification *(17)*. We consider overlapping introns as “alleles” of the same intronic part (Fig. 1c) and calculate the read coverages of the allele “haplotypes” between introns and nucleotide variants. In this way, we can incorporate splicing isoforms in the mutual information calculation to identify allele-specific splicing.

It should be noted that scAllele is a stand-alone tool and only requires a bam file to conduct variant calling and linkage detection. However, pre-processing of the bam file is recommended to achieve optimal results (Supp. Fig. 1).

### Evaluation of variant calls in GM12878 and iPSC cells

We evaluated the variant-calling function of scAllele using scRNA-seq (Smart-seq2) of GM12878 cells and iPSC cells from 3 individuals *(18)*. These individuals were carefully genotyped by the GIAB *(19)* and 1000 Genomes projects *(20)*, thus providing a “ground truth” for method evaluation. We compared the performance of scAllele to those of three other popular variant callers: Freebayes (v.1.3.4), GATK (v.4.2.0.0) and Platypus (v.1.0) *(8, 9, 21)*. The performance evaluation followed previously published guidelines *(22)* with some modifications to accommodate RNA variants (see Methods). For GM12878, we used three benchmark datasets: GIAB’s list of all genetic variants, GIAB’s list of high-confidence genetic variants and the variant calls based on long-read DNA sequencing (Oxford Nanopore) *(23)*.

For each data set and each method, we calculated the true positive counts at specific false positive cutoffs for microindels and SNPs respectively (Fig. 2a, Supp. Fig. 2a). Overall, scAllele achieved the best performance for microindels among all methods, and its performance on SNPs was on par with the others. The strength of scAllele in microindel identification is notable as these variants are known for their challenging detection *(24)*. Furthermore, scAllele also demonstrated superior performance in capturing microindels in “difficult regions” (Fig. 2b, Supp. Fig. 2b). These difficult regions were defined by GIAB *(19)* as the union of regions with low mappability, high GC-content, low complexity or presence of repeats, and segment duplication among others.

**Figure 2.**
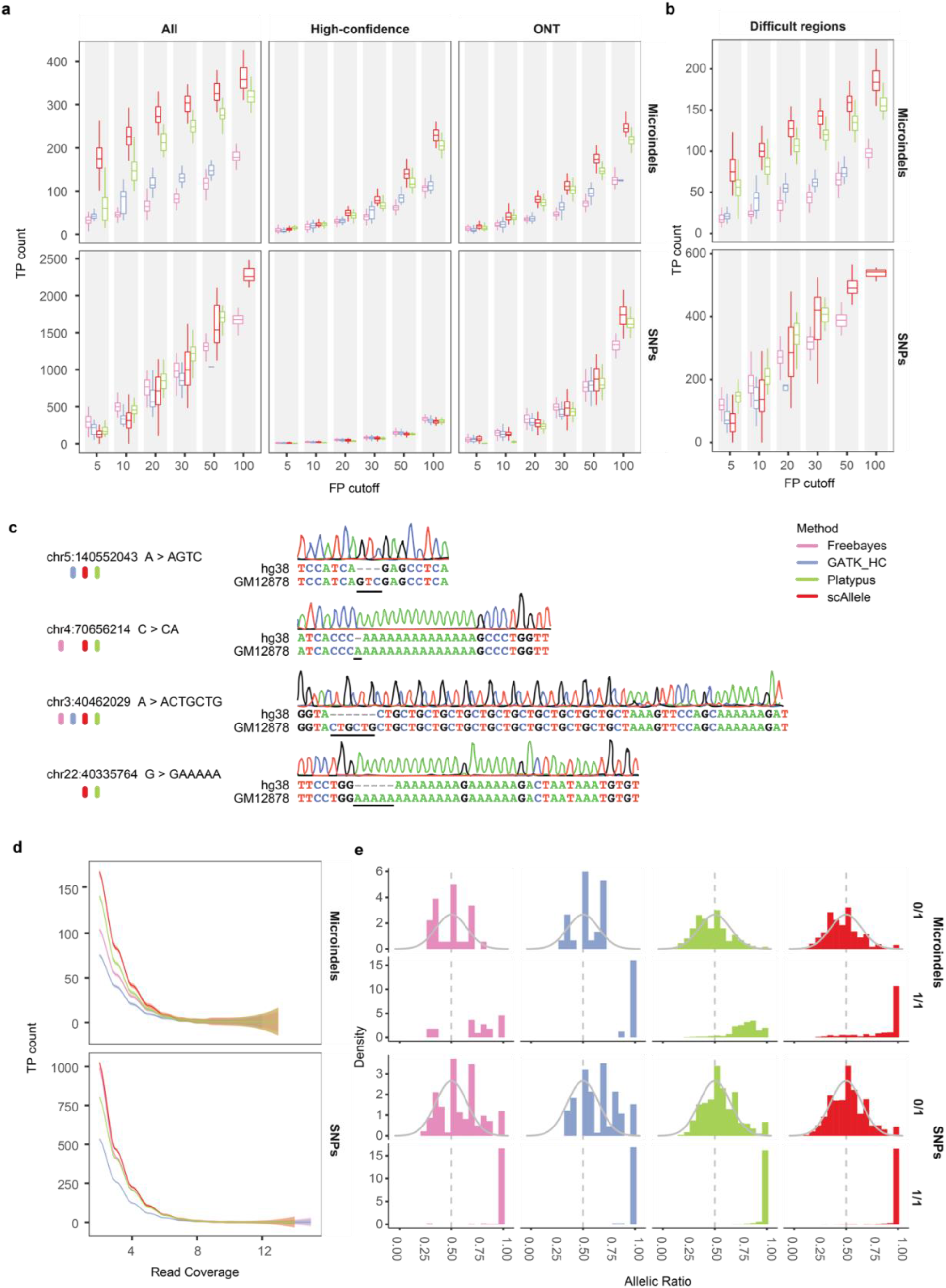
Performance of scAllele in variant calling of scRNA-seq data of GM12878 cells. a. True positive (TP) counts of four variant callers were evaluated at different false positive (FP) cutoffs. Performance for microindels (top) and SNPs (bottom) is shown separately. Three sets of ground truth genetic variants were used: all of GIAB-reported small variants (all), GIAB’s high-confidence variants (high-confidence) and variants called via Nanopore sequencing by Karst et al. (ONT). b. Performance in ‘difficult regions’ defined by GIAB. c. Experimental validation of novel microindels via Sanger sequencing. The bars underneath the variant coordinates indicate the methods that detected the variants. d. True positive count of four variant callers (at maximum F1 and specificity > 0.9) across different read coverages. The curves represent averages of all cells. e Allelic ratios of true positive variants segregated by their true genotype. 0/1: heterozygous; 1/1: homozygous variant allele. Gray curve is shown for reference purpose only (normal distribution of mean = 0.5 and std. dev = 0.15).

It should be noted that although the above cells have been analyzed by the GIAB and 1000 Genomes projects, their genotype calls may still miss some true positives. As examples, we experimentally confirmed 4 microindels categorized as false positives according to the “ground truth” (Fig. 2c, Supp. Fig. 3). The 4 microindels were identified by scAllele and Platypus (2 by GATK, 3 by Freebayes). Thus, the above performance of scAllele (and the other methods) may be a conservative estimation.

One of the hallmarks of scRNA-seq is the limited read coverage per gene. Thus, it is highly desirable to develop variant callers with superior performance at low read coverage. scAllele meets this demand and demonstrates a performance gain relative to the other methods in lowly covered variants (Fig. 2d, Supp. Fig. 2c). Indeed, about 95% of the ground truth variants were covered by less than 5 reads in each data set. Thus, scAllele affords a unique advantage for scRNA-seq data.

Unique to RNA-seq, the allelic read counts of genetic variants reflect their allelic expression levels. Thus, in addition to variant calling, it is necessary to accurately estimate the allelic quantification of each variant. To test the performance of scAllele in this regard, we segregated the ground truth variants into heterozygous and homozygous groups. The heterozygous variants are expected to exhibit an approximately normal distribution in their ALT allelic ratios (variant allele read number/total read number), centered around 0.5 *(13)*. For homozygous variants, the allelic ratios are expected to be 1. As shown in Fig. 2e (and Supp. Fig. 2d), the results of scAllele largely followed these expectations for both microindels and SNPs. In contrast, other methods resulted in flawed distributions in at least one of the above aspects.

Overall, the above evaluations support the superior performance of scAllele for scRNA-seq variant analysis, especially in handling microindels, an aspect that is much more challenging compared to the most-often tackled SNV identification.

### Linkage calculation between variants

In addition to variant calling, scAllele enables read-level allelic linkage analysis. Such analysis is not possible with other variant callers as read-level information is not extracted. In scAllele, the degree of allelic linkage is quantified as the mutual information (MI) between two types of variants: nucleotide variants and alternatively spliced junctions (Fig. 1). This metric is expected to require a relatively high number of reads harboring both types of variants. To achieve an understanding of the read coverage requirements, we first calculated the MI between pairs of known genetic variants in the GM12878 and iPSC data used in the last section. As expected, the MI of these variant pairs in the RNA is generally high, regardless of read coverage, reflecting the associated DNA haplotypes (Fig. 3a).

**Figure 3.**
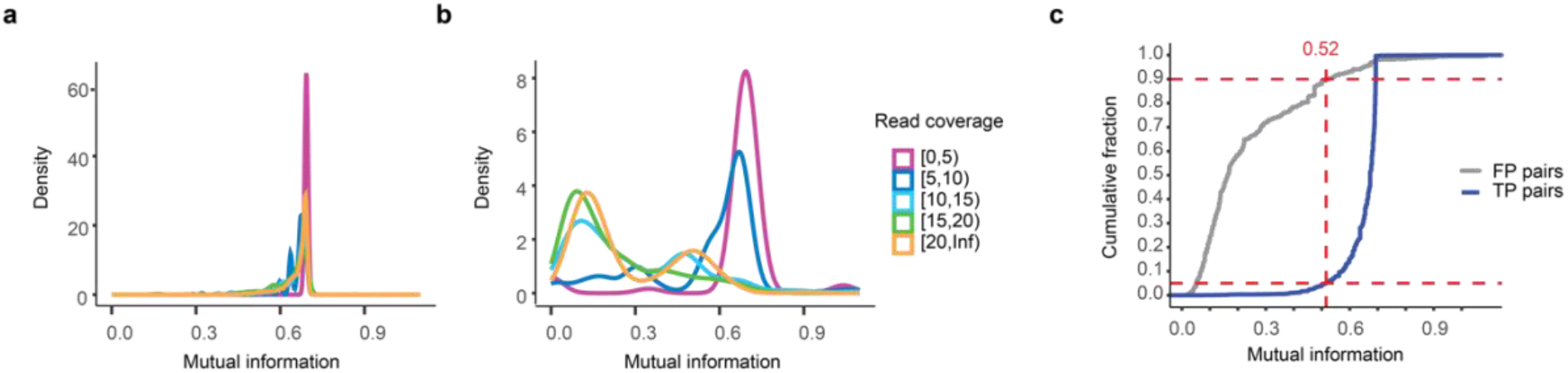
Linkage calculation between pairs of variants. a. Mutual information (MI) distribution (natural log-based) of pairs of true genetic variants (namely, true positives (TPs)) from the GM12878 and iPSC scRNA-seq segregated by the number of reads covering the pair. Most of these pairs have values close to the theoretical maximum for two alleles (ln(2) = 0.693) regardless of coverage. b. MI distribution of pairs of variants where at least one is not a known genetic variant (here referred to as false positive (FP) genetic variants). c. Cumulative distribution of MI of TP and FP pairs with a minimum read coverage of 10. The MI cutoff of 0.52 was selected as the minimum value for significant linkage between variants. This cutoff rejects 90% FP pairs and 5% TP pairs (dashed lines).

As a comparison, we also calculated the MI between pairs of nucleotide variants where at least one variant was not a known genetic variant (Fig. 3b). Since the cell lines have been well-genotyped, we assume all unknown variants observed in the RNA-seq reads are RNA editing sites or sequencing errors. The MI of these variant pairs is expected to be low in general *(17)*, unless rare allele-specific RNA editing exists. This expectation of low MI was met at relatively high read coverage (>10). However, at lower read coverage, the MI is inflated due to the low number of transcripts used for its calculation. Thus, it is necessary to impose a minimum read coverage requirement for MI calculation. In this study, we set this cutoff to be 10 based on the above results. Additionally, we required a minimum MI of 0.52 to call significant linkage events, as 95% of the known genetic variant pairs (with ≥10 reads) had an MI of 0.52 or greater, and 90% of the unknown variant pairs (with ≥10 reads) failed this MI cutoff (Fig. 3c). The read coverage and MI cutoffs can be altered by the user in scAllele.

### scAllele unveils nucleotide variants and allele-specific splicing events in lung cancer cells

Next, we applied scAllele to scRNA-seq data of lung cancer (Smart-seq2) *(25)*. We focused on cancer cells and their normal counterparts, epithelial cells, in tumor and matched normal samples of two patients (TH179 and TH238; n = 574 cells). We first carried out variant calling in each cell. An SNV or microindel was retained if it was detected in at least 3 cells. Furthermore, we compared the presence of the variants in normal epithelial or cancer cells. A variant was defined as cancer-enriched if it was not detected in normal cells or its presence is significantly more frequent in cancer compared to normal cells (BH-corrected p < 0.1, Methods). Otherwise, the variant was labeled as a common variant to cancer and normal cells. As a sanity check, we note that no variant was found to be enriched in normal cells relative to cancer cells.

As shown in Fig. 4a, >15,000 variants were identified in each patient, with the majority being SNVs. Most SNVs are annotated variants in the dbSNP (b151) or Cosmic (human cancer mutations) database. Importantly, Cosmic variants constitute a larger fraction among cancer-enriched variants compared to the common variants (p < 1e-14 in both patients, Chi-Squared test). Among the unannotated (i.e., novel) SNVs, some may be novel genetic variants and others may reflect RNA editing events. Indeed, a large fraction (74% in TH179, 76% in TH238) of the novel SNVs corresponded to A-to-G or C-to-T RNA editing types. A relatively large fraction of microindels was not annotated in either database, likely reflecting our incomplete knowledge of this type of variants. Interestingly, cancer-enriched SNVs were more often located in coding exons, less often in introns, compared with common SNVs (Fig. 4b). In contrast, the fraction of microindels in coding exons is approximately the same for the two categories of variants. Nonetheless, one patient (TH179) showed enrichment of cancer-enriched microindels in 3’ UTRs (depletion in introns), relative to the common microindels.

**Figure 4.**
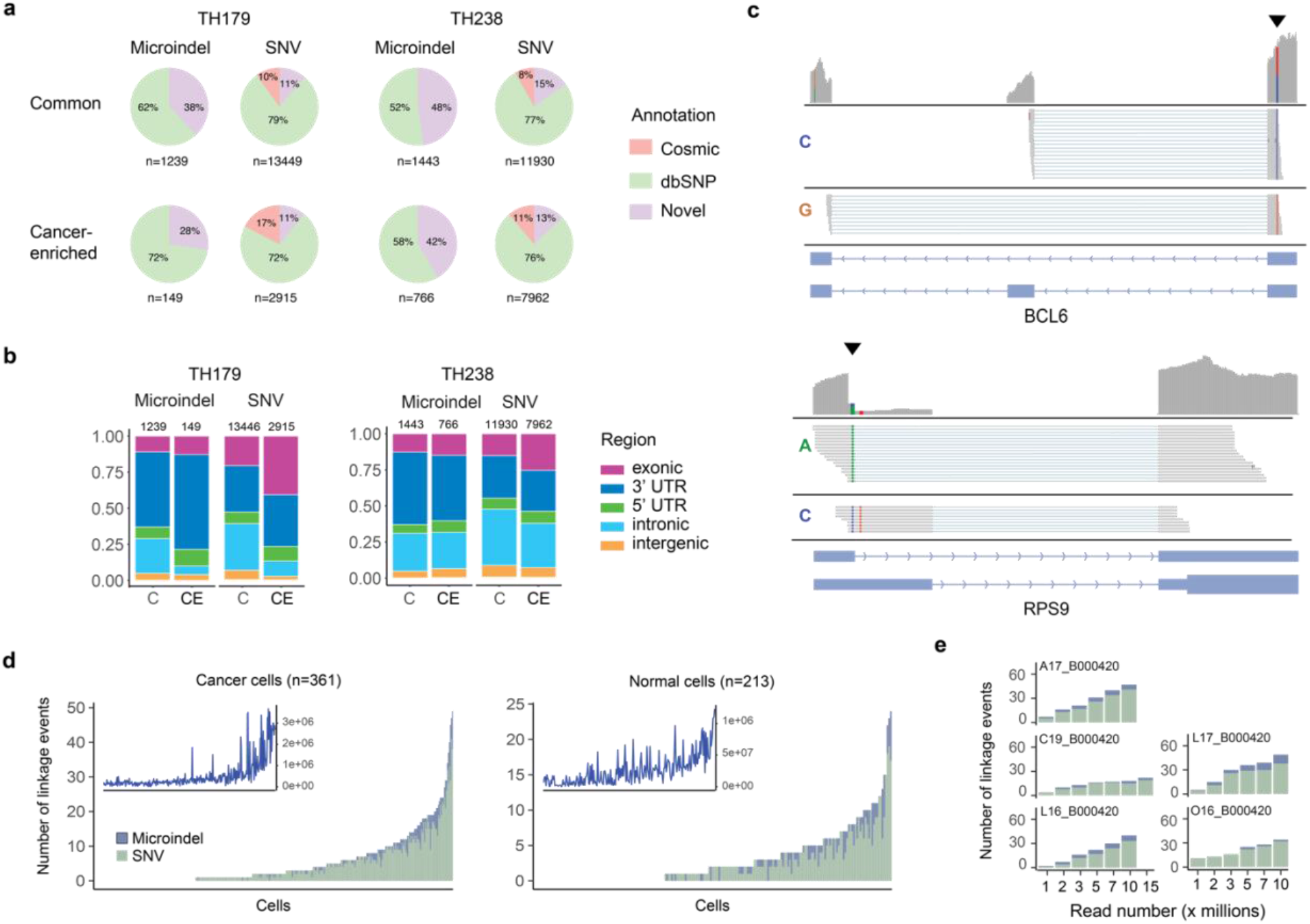
Summary of variants and linkage events detected by scAllele in the Lung Cancer scRNA-seq dataset. a. Variants identified in each patient (≥3 cells). Common: variants common to cancer and normal cells. Cancer-enriched: variants enriched in cancer cells (see Text). Novel: variants not annotated in dbSNP or Cosmic. Variants labelled as “Cosmic” may also present in dbSNP. b. Distribution of variants in a in different types of genomic regions. C: common. CE: cancer-enriched. c. IGV view of two example allele-specific splicing events. The location of the variant is denoted by the black arrowhead. The reads overlapping each variant are segregated according to the allele they harbor (indicated by the base color and the base label on the left). d. Number of linkage events identified in cancer or normal cells (union of cells from the two patients). The cells are ranked by their total number of events. Events associated with microindels or SNVs are shown in different colors. The insets show the total number of junction reads in the scRNA-seq data of the cells sorted in the same order as the main panel. e. Number of linkage events identified in 5 deeply sequenced cells at different down sampled total read coverage.

Following variant calling, we identified the allele-specific splicing events in each cell. As examples, Fig. 4c shows two significant linkage events. In these cases, the SNPs demonstrated strong allelic linkage with alternative splicing patterns (exon skipping and alternative 5’ splice site, respectively). Across cancer and normal cells, the number of allele-specific splicing events varied greatly, ranging from 0 to 49 events (Fig. 4d), most of which involved SNVs. This number correlated approximately with the number of spliced-junction reads present in each cell (Fig. 4d insets). Based on down-sampling of a few deeply sequenced cells, we observed that 1M total reads can enable identification of up to 11 events per cell (Fig. 4e). In some cells, the number of events plateaued at around 5M reads. Thus, to afford power for splicing analysis, a relatively large number of scRNA-seq reads is needed.

### Cancer and normal cells exhibit unique and differential linkage events

Next, we asked whether cancer and normal cells harbor different allele-specific splicing events. Among all events identified in this study, 27 were observed in both cancer and normal cells, whereas more events were exclusive to one of the two classes of cells (Fig. 5a). In general, most events were observed in a small number of cells (< 5), possibly due to read coverage limitations.

**Figure 5.**
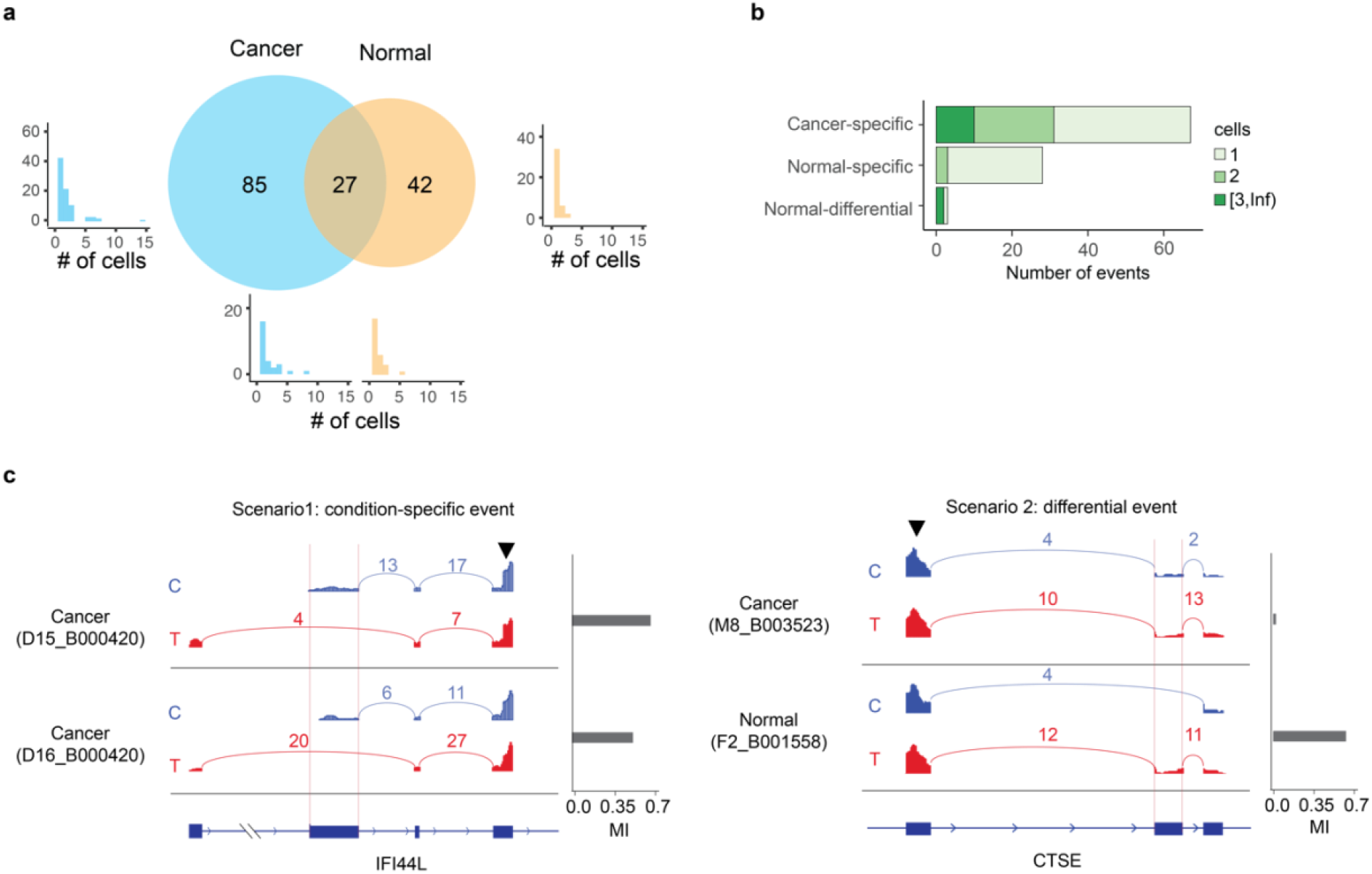
Comparison of allele-specific splicing events in cancer and normal cells. a. Number of events shared by cancer and normal cells, or exclusive to one of the two classes. Histograms of the number of cells harboring each type of events are also shown for each set and the intersect. b. Number of events categorized as cancer-specific, normal-specific, or differential (see Text for details). The number of cells harboring the events are shown by the shade of the bar. c. Examples of allele-specific splicing events (red vertical lines) from condition-specific events (left) and differential event (right). The sashimi plots are split by the allele harbored in the reads (indicated by color and base label). The read counts are reported for each junction and the mutual information between the variant and the splicing event is shown by the bars on the right. Note that only reads harboring the variant (black arrowhead) are shown.

To identify differential linkage events between cancer and normal cells, we focused on two scenarios. In the first scenario, the variants were not present/testable in the normal cells but had significant linkage in the cancer cells, or vice versa (labeled as “cancer-specific” or “normal-specific”, Fig. 5b, Table S1). For this scenario, we observed 67 events that were cancer-specific and a smaller number of events (29) that were normal-specific. Notably, 11 cancer-specific events were observed in at least 3 cancer cells, whereas no normal-specific events exceeded this level of prevalence (Fig. 5b). Figure 5c (left) shows an example in the gene IFI44L (interferon-induced protein 44-like), a type I interferon stimulated gene with a role in host antiviral response *(26)*. The allele-specific splicing event and the associated variant were only observed in cancer cells.

The second scenario includes variants present and testable for splicing linkage in both cancer and normal cells, but significant linkage was detected with higher prevalence (p-value < 0.05 Fisher’s exact test) in one cell class than the other. For this scenario, we observed 2 events, both of which occurred in higher proportion in the normal than cancer cells (Normal-differential, Fig. 5b, Table S1). These events may reflect a loss of function of the variants in the cancer cells. Figure 5c (right) shows an example of such an event in the CTSE gene, where the C allele of the variant is linked to skipping of the middle exon, whereas the T allele is associated with exon inclusion. This linkage was only observed in normal cells, but not cancer cells (despite the presence of the variant and adequate read coverage in cancer cells). Notably, the gene CTSE encodes for Cathepsin E, an aspartic protease with a vital role in protein degradation, bioactive protein generation, antigen processing and presentation *(27)*.

## Discussion

scRNA-seq affords unprecedented views of single cell transcriptomes. Similar to bulk RNA-seq, scRNA-seq provides information on the single-nucleotide level. However, identification of nucleotide variants in scRNA-seq is challenging due to the limited read coverage per cell. Here, we present scAllele, a versatile tool that not only enables variant calling in single cells but also uncovers allele-specific RNA processing events.

We showed that scAllele outperforms other popular methods in variant calling, especially for microindels, the class of variants that are less well characterized than SNVs. Built upon local reassembly, scAllele refines read alignments and corrects possible misalignments in each read, thus enhancing variant detection accuracy per read. Additionally, scAllele uses a GLM model to detect high confidence variants. These features together confer an advantage in performance given low sequencing depth.

The read-level variant calling by scAllele enables another advantage, that is, facilitating a detailed view of the allelic bias linked to alternative RNA isoforms. In this work, we focused on allele-specific splicing patterns. A similar approach can be extended to examine other aspects of RNA expression, such as alternative polyadenylation. We note that this type of analysis requires a relatively high read coverage per event, as it simultaneously quantifies alternative alleles of nucleotide variants, alternative RNA isoforms and their combined linkage patterns. We showed that the number of such events increased with higher scRNA-seq sequencing depth, indicating that RNA representation in scRNA-seq was not saturated at lower depth, such as 1M reads. With the continued drop in sequencing cost, we expect to see wide applications of allele-specific and alternative RNA isoform analyses, such as those enabled by scAllele.

We applied scAllele to a lung cancer scRNA-seq dataset (with matched controls). Our analysis identified a large number of nucleotide variants, many of which had enriched presence in cancer cells. Compared to variants common to both normal and cancer cells, cancer-enriched variants were more often cataloged in Cosmic, supporting the validity of the scAllele variant calls. Additionally, cancer-enriched variants were more often located in coding regions, which suggests a potential role in altering protein sequences or producing neoantigens. Although microindels are not as abundant as SNVs, they may also have critical roles in human diseases *(28)*, an area remains under-explored. In general, compared to SNVs, we observed a relative enrichment of microindels in 3’ UTRs. Given the existence of numerous regulatory elements in the 3’ UTRs *(29)*, microindels in these regions may alter many processes, such as mRNA stability, translation, or mRNA localization, which should be investigated in the future.

In the cancer and normal epithelial cells, we identified more than 150 allele-specific splicing events. We further categorized these events based on their relative prevalence in cancer or normal cells. Even though most events were observed in a small number of cells, likely due to low read coverage in single cells, such categorization provides an approximate overview of their relative enrichments. Among these events, many have important relevance to cancer, such as those in the CTSE and IFI44L genes in Fig. 5c. Our results suggest that scRNA-seq data possess useful information to uncover important alternative splicing events, linking genotypes to this molecular phenotype.

In summary, scAllele offers a unique approach to maximize the information extracted from scRNA-seq data sets. With the emergence of scRNA-seq data from a large spectrum of samples, scAllele will lead to a granular view of the genetic landscape of each cell and the potential genetic drivers of gene expression complexity.

## Methods

### scAllele: detailed outline

To scan variants in the entire transcriptome, we grouped the sequencing reads into Read Clusters (RC). An RC is made up of a group of overlapping reads. It should be noted that the entire sequence of each read was included, not only the segment that overlaps the RC. Multi-mapped, chimeric, and low mapping-quality reads were removed as well as reads with ≥ 5 soft-clipped bases or trailing homopolymers (n ≥ 15). An additional “reference read” was included as part of the RC. This read contains the genomic reference sequence of the entire range spanned by the RC.

In each RC, the reads were decomposed into overlapping k-mers (k-1 overlap) which are the nodes of the de Bruijn Graph (dBG). The edges represent consecutive nodes (i.e., two k-mers overlapping by k-1) in the reads. Every edge was labeled with the name of the reads that contained this consecutive pair of k-mers and the position in the read where the k-mers were located.

The graph was then processed by compacting and removing certain nodes. Walks on the graph that contain consecutive nodes of *in-degree = 1* and *out-degree = 1* can be merged into a single node that contains a sequence length of *k + n – 1* where *n* is the number of nodes being merged. In addition, subsequences in the “reference read” that did not overlap with other reads (which are usually intronic segments) were also compressed. This step greatly simplifies the graph because the intronic regions are generally several thousand bases long, much longer than the average RC. Other nodes were removed from the graph if they did not provide useful information. For example, we defined the actual start and end of the RC as the first and last nodes that originated from the “reference read”. By definition, these nodes have *in-degree = 0* and *out-degree = 0* respectively. Additional nodes that complied with the degree requirement but did not originate from the “reference read” were labeled as alternative starts and ends. These alternative starts and ends also represent differences among read sequences. However, since they do not form a bubble, it’s not possible to infer the variant causing such difference.

Subsequently, scAllele inferred the walk on the graph that matched the original sequence of each read. These walks were named “read walks”. Because some nodes were removed or merged in the previous step, this walk is not necessarily the same sequence of nodes obtained from the initial read decomposition. As a result, many of the original reads were matched by the same “read walk”, reducing the number of distinct reads to process.

In the compacted/cleaned dBG, we identified the bubble structures by locating the source nodes, the sink nodes and the walks connecting them via depth-first search (DFS) of the graph. These structures represent variants, and with the DFS we can identify which specific source node, sink node, and connecting walk correspond to each allele. This information was then used to identify the variants and their alleles present on each “read walk”. In the case of highly interconnected/cyclic graphs (due to existence of repeats or low complexity regions), this assignment was aided by a Dijkstra-like algorithm which identifies the most likely set of variants on a “read walk” by minimizing the editing distance between the read walk sequence and the reference sequence. More specifically, first, all the end-to-end “read walks” were identified.

Then, by calculating the cumulative edit distance at every node and traversing the graph through different walks, we can select the best walk. Note that introns were also considered a type of variants in these intermediate steps and were processed in the same way as the nucleotide variants. However, we did not assign an edit distance to them.

The variants were further processed by normalizing, left aligning and atomizing them. Different features were collected for each variant including read counts for each allele, base qualities, read positions, and count of tandem repeats flanking the variant. An additional feature, namely haplotype-fitting was calculated using the entire set of reads and variants from each RC. These features were then used to score the quality of the variant (see Variant Scoring). At this point, the variant-calling step was complete. Since scAllele identifies variants at the read level, this information was stored in memory for subsequent mutual information analysis, based on which the linkage between nucleotide variants and splicing isoforms was calculated (see linkage analysis).

### Variant Scoring

We trained a generalized linear model (GLM) using “ground truth” genetic variants and various features obtained from the main algorithm of scAllele. The features included the variant’s ALT allelic ratio (AB), the number of tandem repeats neighboring the variant *(TandemRep)*, sequencing error rate (SER), median base quality in the variant’s proximity, read position, and the haplotype-fitting, as detailed below.

Low ALT allelic ratio is often indicative of a false positive variant, likely due to existence of sequencing errors. In fact, we can calculate the probability of observing an allelic ratio (AB) if it is resulted from a sequencing error using the binomial distribution.

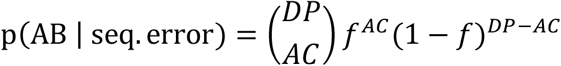

where *AC* is the ALT allele counts, and *DP* is the total read count. The value of *f* is the probability of error which in most cases corresponds to the SER. We used 0.01 for this variable, which is the maximum error rate for the Illumina sequencing platforms *(30)*. In tandem repeats, however, the probability of error is expected to increase due to the propensity of PCR slippage. We then defined *f* as follows:

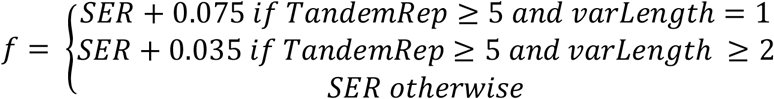

These values are approximations of the empirical estimation of stutter noise made in lobSTR *(31)*. The stutter noise was found to be a function of the variant length *(varLength)* and the number of tandem repeats *(TandemRep)*. The probability p(AB|seq.error) was used as an additional feature in the GLM, and served as an interaction term between *AB* and *TandemRep*.

Another feature in the GLM was derived from base quality scores. In case of SNVs, we simply used the base quality at the mismatch position. For microindels, we used the median base quality in the neighboring region of the variant (+/- 7 bases) since the original position of the microindel is, in many cases, ambiguous. In addition, the median read position was used as a variable in the GLM because the 3’ ends of the reads tend to have lower base quality, also considering the fact that the bubble structures in the dBG are less reliable if they only use the ends of the “read walks”.

Next, scAllele calculated another metric called “haplotype-fitting”. This refers to the ability to cluster the variant alleles into two potential haplotypes based on their colocalization in the reads. We clarify that we do not aim to infer the actual haplotype since RNA-seq data is not ideal for this task. This step simply checks for multi-allelic variants and allele combinations that result in more than two haplotypes. For this step, we discarded potential RNA editing sites and we perform the clustering at the RC level which, most of the time, matches exonic coordinates. In this way, the haplotype is not confounded by non-genetic variants or allele-specific splicing.

The regression of the GLM was performed using the scikit-learn package *(32)* from Python. The training data consist of genetic variants identified in scRNA-seq data originated from the GM12878 cell line. We used the ground truth from GIAB *(19)* and trained the model on a subset of the dataset used in the “*Evaluation of variant calls in GM12878 and iPSC cells”* section. It’s important to note that the training data was not used to derive the performance results in that section. We trained a separate model for SNPs, insertions, and deletions respectively. The data was shuffled and split 25 times for cross-validation.

Lastly, we sought to define a “quality score” for the scAllele variant call. First, we consider the log-likelihood of the GLM regression as a regression score:

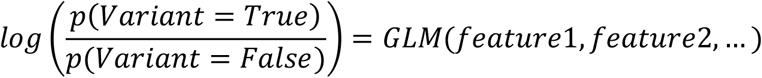

Meanwhile, the quality score (QUAL) of a standard VCF file format (specified by VCFtools, v.4.2) is a Phred-scaled form:

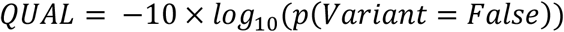

Thus, a GLM regression score of zero corresponds to QUAL = 3.01, which represents equal probability of a variant being true or false. Based on the benchmark evaluation, we observed that the F1 score was usually maximized at regression scores between 1 and 2 (10.4 ≤ QUAL ≤ 20.0). Thus, in scAllele, the default score format is QUAL with a cutoff of 10. However, the user can choose to use regress scores to define quality of variant calls and the score cutoff can also be defined by the user.

### Linkage analysis

scAllele detects variants at the read level allowing for allelic linkage detection. For every RC, all the reads that overlap a variant position were collected with their corresponding allele recorded (REF or ALT). Reads from different RCs were pooled together after scanning an entire chromosome. In paired-end data, a RC may not contain both mates of the pair. Thus, by merging reads from different RCs we can increase the number of potential linkages.

For every pair of variants that were less than 100 kb apart, scAllele retrieved the reads that overlapped both variants. Using these reads, one array per variant was constructed containing the allele information of the variant. We used Python’s scikit-learn package *(32)* to calculate the mutual information between these two arrays.

For the calculation of linkage between nucleotide variants and splicing isoforms, scAllele first grouped overlapping introns into “intronic part” (Fig. 1a). Such overlapping introns were considered as “alleles” of the intronic part. In this way, the mutual information between an intronic part and a nucleotide variant can be calculated similarly as for a pair of nucleotide variants.

Based on our analysis in the section “*Linkage calculation between variants”*, we required a minimum of 10 common reads for a pair of variants. Additionally, a minimum mutual information of 0.52 was required to define a significant linkage.

### scRNA-seq processing and mapping

Raw scRNA-seq fastq files from the GM12878 cell line were retrieved from ENCODE (Accession: ENCSR000AIZ). These replicates were deeply sequenced (about 30 million reads). Thus, we downsampled each replicate into 30 alignment files with roughly 1 million reads each. The goal was to resemble a shallowly sequenced sample to test our method on low coverage data.

The scRNA-seq data from iPSCs, corresponding to individuals NA19098, NA19101, and NA19239, were obtained from NCBI (Accession number: GSE77288) (http://www.ncbi.nlm.nih.gov/geo/query/acc.cgi?acc=GSE77288)

The lung cancer *(25)* dataset was also downloaded from NCBI (BioProject PRJNA591860) (www.ncbi.nlm.nih.gov/bioproject/28889).

Raw reads from all samples were pre-processed using fastqc (v.0.11.7) *(33)* to check for adapter content and over-represented sequences. If present, these sequences were removed using cutadapt (v.1.9) *(34)*. 3’end of reads with low base quality were also trimmed using sickle (v.1.33)*(35)*. The reads were aligned using two-pass STAR alignment (v.2.7.0c) *(36)*. Finally, we marked PCR duplicates using the tool MarkDuplicates from Picard Tools (v.2.25.2) *(37)* (Supp. Fig. 1).

### Evaluation of multiple variant callers

Variants were called by three other tools: Platypus, GATK-HaplotypeCaller and Freebayes. The recommended pre-processing steps and parameters were used. For Platypus, the variant call was performed after processing the alignment file with Oppossum. For GATK, the variant calls were carried out following the “best practices” steps for RNA variant calling. For Freebayes, the variant call was performed with default parameters. These variants were then filtered to remove predicted variants with alternative (ALT) allele read count < 2 and A-to-G, C-to-T mismatches as they may represent RNA editing sites. Variants with labels indicating low quality were also removed.

We evaluated the performance of the variant callers using the vcfeval function from rtg tools (v.3.12) *(38)* according to the benchmarking standards reported previously *(22)*, with the following parameters:

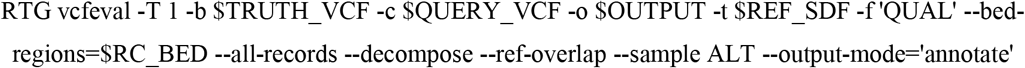

The variable RC_BED is a bed file containing all the genomic regions covered by at least one read. We employ it to reduce the running time of the software. The option “--sample ALT” was used to skip genotype matching and is more appropriate for RNA data.

The performance metric used in this study was true positive count at fixed false positive thresholds. Given that the ground truth genetic variants were obtained from WGS, the sensitivity of variant calling in RNA-seq is in principle restricted to the number of genetic variants that are transcribed and sequenced in RNA. Hence, sensitivity, true-positive rate and F1 scores won’t accurately reflect the performance of variant callers in RNA-seq.

For the evaluation of variant calling in difficult regions, we used the union of bed files containing difficult regions for variant calling from GIAB which merges regions of low-mappability, high GC-content, segment duplication, low complexity, functional regions, and other difficult regions.

### Detection of cancer-enriched variants and annotation

We applied scAllele to the lung cancer dataset from Maynard et al *(25)* and selected cancer and normal epithelial cells corresponding to two individuals (TH238, TH179) where both cancer and normal tissue biopsies were obtained. Then, we focused on variants present in at least three cells of an individual. We calculated the prevalence of each variant across cells. Using the hypergeometric test, we evaluated the enrichment of each variant in cancer cells compared to normal. The p-value obtained was then corrected for multiple testing using the BH method. We defined a given variant as cancer-enriched if the BH corrected p-value is ≤ 0.1 or if the variant is not present in any normal cell.

We further overlapped the variants with the COSMIC (cancer.sanger.ac.uk) *(39)* and, subsequently, dbSNP (b151) *(40)* databases. From the COSMIC annotation, we only selected variants that were confirmed to be somatic and were found in lung tissue. A variant was labeled “novel” if it is not present in either database.

### Detection of differential linkage events

To detect differential linkage, we selected common linkage events (same nucleotide variant and same introns) between the cells of the same individual. We then classified them into four categories explained by the two proposed scenarios. For scenario 1, we selected linkage events that were present in cancer cells, but not in normal cells, or vice versa. These events were grouped into the categories “cancer-specific” and “normal-specific”, respectively. For scenario 2, we selected linkage events that were present in both types of cells, but significantly more prevalent in one compared to the other. These events were grouped into the categories “cancer-differential” and “normal-differential”. To detect differential prevalence, we used the Fisher’s Exact test using the number of cells with the linkage event and the number of cells that were testable for linkage in each group of cells.

## Code availability

The scAllele software is available in our github repository (https://github.com/gxiaolab/scAllele/). The scripts used for the analyses in this work are available at: https://github.com/gxiaolab/scAllele/Manuscript

## Data availability

Variant calls and linkage events from the GM12878, IPSC cells for individuals NA19098, NA19101 and NA19239, and the lung cancer cells are available in our github repository: https://github.com/gxiaolab/scAllele/data

## Acknowledgements

We thank members of the Xiao laboratory for helpful discussions and comments on this work. We appreciate the helpful discussions with Dr. Serghei Mangul. This work was supported in part by grants from the National Institutes of Health (U01HG009417, R01AG056476 to X.X.) and the Jonsson Comprehensive Cancer Center at UCLA.

