## Supplementary material for "scAllele: a versatile tool for the detection and analysis of variants in scRNA-seq": Supp. Fig. 1

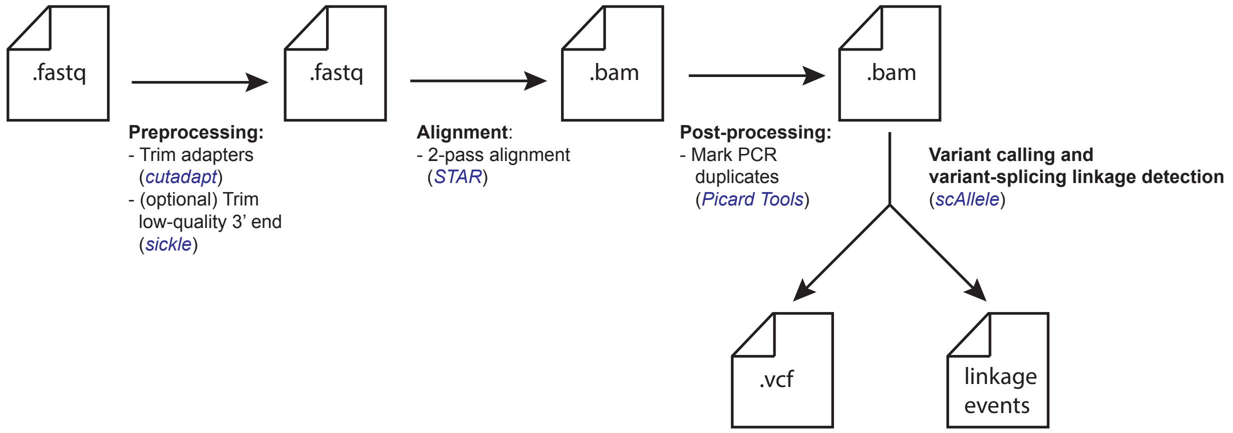


**Supplemental Figure 1.** Recommended step-by-step pipeline for variant-calling and variant-splicing linkage detection. Reads from raw fastq files were processed to trim adapters and low quality 3’ ends (names of the tools shown in blue). These reads were then aligned by STAR using the 2-pass alignment option. This option allows for accurate mapping of splice junctions which is important for accurate identification of variant-splicing linkage. After alignment, PCR duplicates were marked using Picard Tools. Duplicated reads amplify the frequency of random errors which can lead to false positive calls. Finally, scAllele calls variants and identifies variant-splicing linkage.

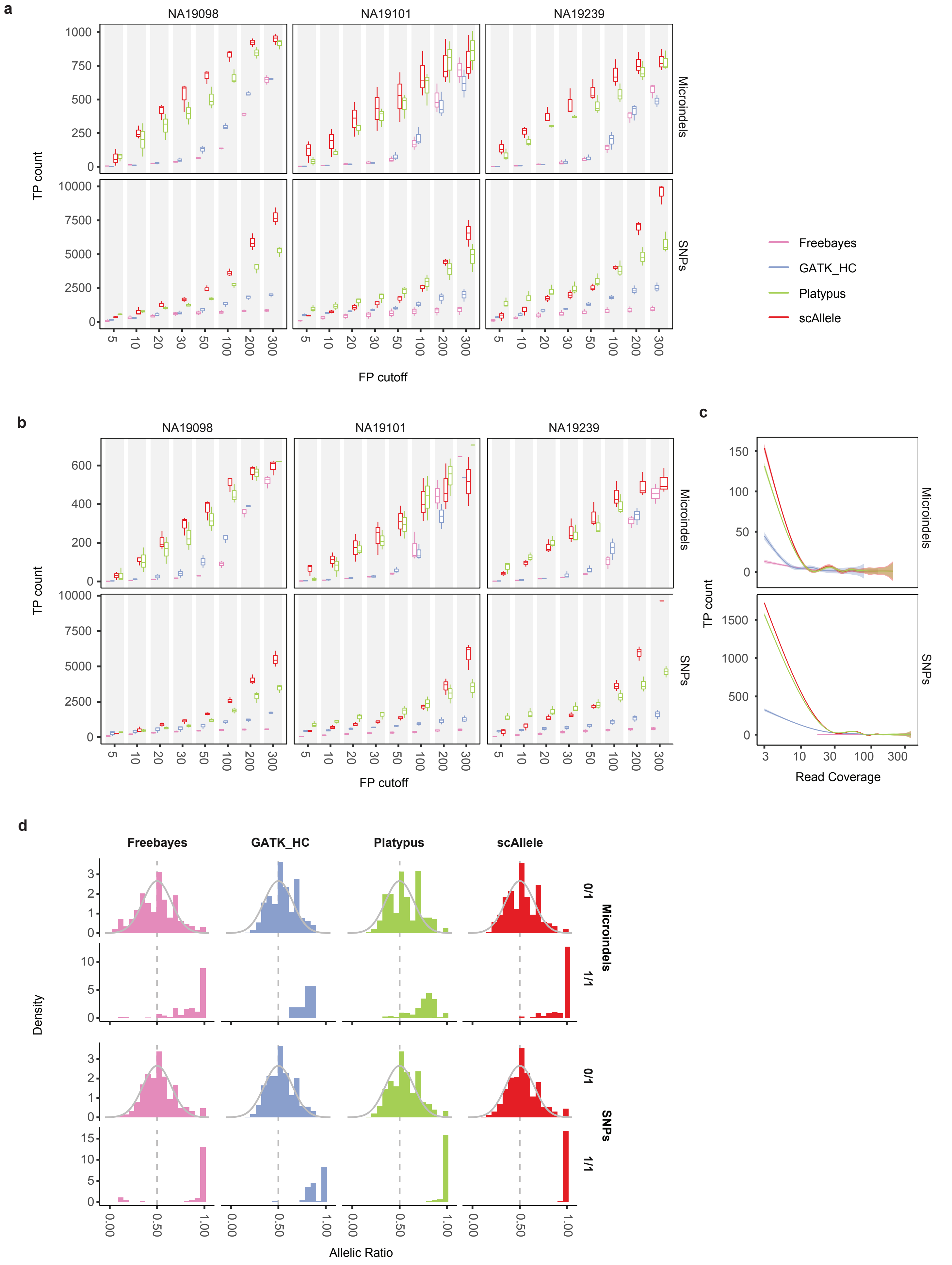


**Supplemental Figure 2.** Performance of scAllele in variant calling of iPSC scRNA-seq of three individuals. **a.** True positive (TP) count of four variant callers were evaluated at different false positive (FP) cutoffs. Performance for microindels (top) and SNPs (bottom) is shown separately. All GIAB-reported small variants are included here. **b.** Performance in 'difficult regions' defined by GIAB. **c.** True positive count of four variant callers (at maximum F1 and specificity > 0.9) across different read coverages. The curves represent averages of all cells. **d** Allelic ratios of true positive variants segregated by their true genotype. 0/1: heterozygous; 1/1: homozygous variant allele. Gray curve is shown for reference purpose only (normal distribution of mean = 0.5 and std. dev = 0.15).

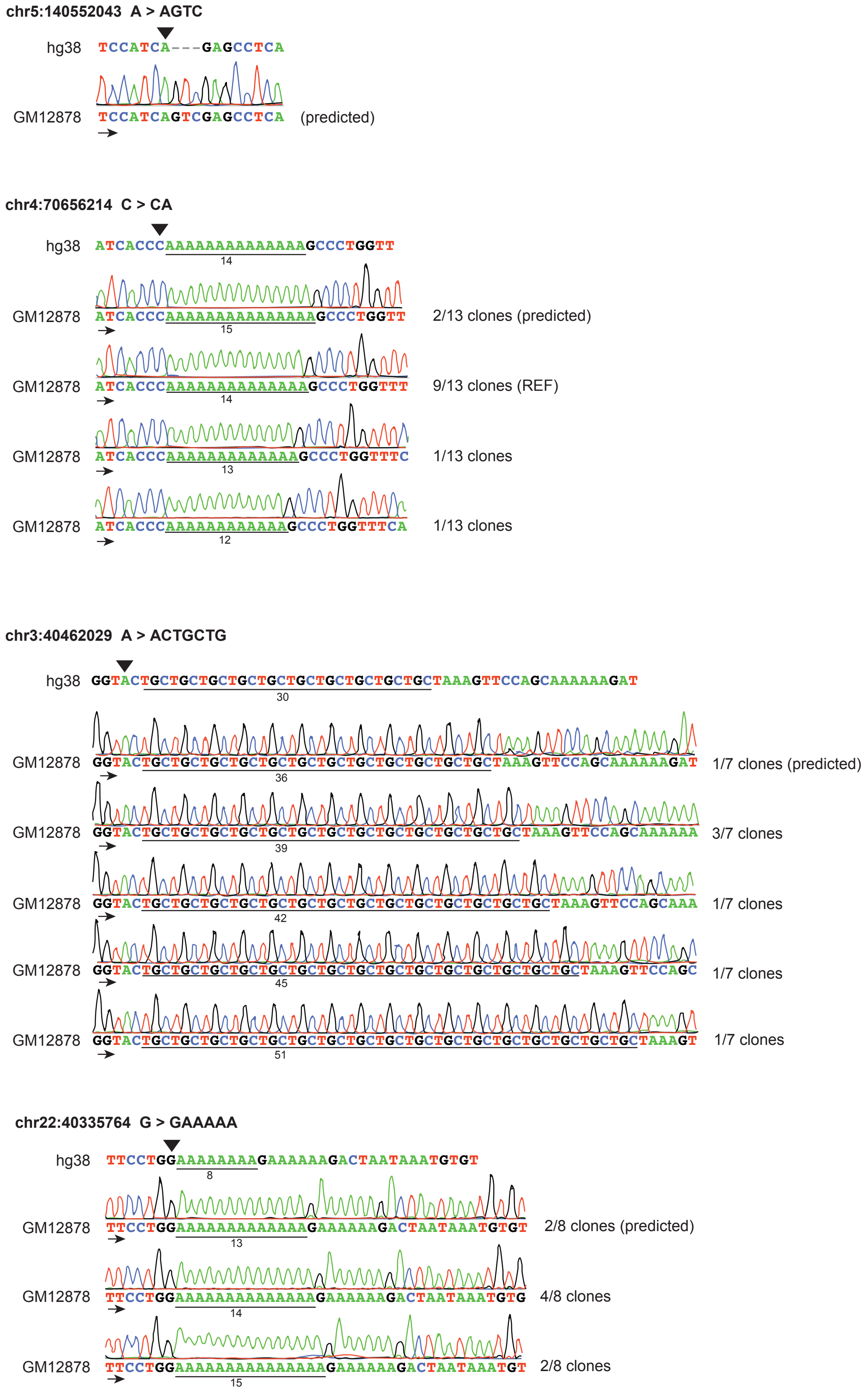


**Supplemental Figure 3**. Experimental validation of novel microindels identified in GM12878 cells. The predicted variant information is shown (chromosome, position, reference allele, and alternative allele). The black triangle indicates the position of the variant in the hg38 reference sequence. The different traces correspond to all the variants detected by Sanger sequencing in GM12878 cells, including the one predicted by the variant callers. The number of clones containing each microindel is also shown. 3 of the 4 tested variants were located adjacent to homopolymers or tandem repeat sequences (which are underlined). The arrow indicates the direction of sequencing.
